# Parallel evolution of gene expression between trophic specialists despite divergent genotypes and morphologies

**DOI:** 10.1101/180190

**Authors:** Joseph A. McGirr, Christopher H. Martin

## Abstract

Parallel evolution of gene expression commonly underlies convergent niche specialization, but parallel changes in expression could also underlie divergent specialization. We investigated divergence in gene expression and whole-genome genetic variation across three sympatric *Cyprinodon* pupfishes endemic to San Salvador Island, Bahamas. This recent radiation consists of a generalist and two derived specialists adapted to novel niches – a ‘scale-eater’ and a ‘snail-eater.’ We sampled total mRNA from all three species at two early developmental stages and compared gene expression with whole-genome genetic differentiation among all three species in 42 resequenced genomes. 80% of genes that were differentially expressed between snail-eaters and generalists were up or downregulated in the same direction between scale-eaters and generalists; however, there were no fixed variants shared between species underlying these parallel changes in expression. Genes showing parallel evolution of expression were enriched for effects on metabolic processes, whereas genes showing divergent expression were enriched for effects on cranial skeleton development and pigment biosynthesis, reflecting the most divergent phenotypes observed between specialist species. Our findings reveal that even divergent niche specialists may exhibit convergent adaptation to higher trophic levels through shared genetic pathways. This counterintuitive result suggests that parallel evolution in gene expression can accompany divergent ecological speciation during adaptive radiation.

**Impact Summary:** Adaptations that result in unique forms of ecological specialization are central to research in evolutionary biology, yet little is known about their molecular foundations. We combined transcriptome sequencing with whole-genome divergence scans to study the molecular evolution of two specialist *Cyprinodon* pupfish species – a ‘scale-eater’ and a ‘snail-eater’ – that rapidly diverged from a sympatric generalist ancestor within the last 10,000 years. While parallel evolution of gene expression driving convergent niche specialization seems common, we present, to our knowledge, the first example of significant parallel changes in expression coinciding with divergent niche specialization. 80% of genes that were differentially expressed between snail-eaters and generalists showed the same direction of expression in scale-eaters relative to generalists. Furthermore, parallel evolution in expression seem to be controlled by unique genetic variants in each specialist species. Genes showing parallel changes in expression were enriched for metabolic processes that may facilitate adaptation to a higher trophic level, while genes showing divergent expression likely shape the striking morphological differences between specialists. These findings contribute to a more nuanced understanding of convergent adaptations that arise during speciation, and highlight how species can evolve similar expression profiles adapted to divergent niches.

## Introduction

Abundant research on the genetic basis of adaptive traits has revealed an overarching pattern in nature – when species are faced with similar selective pressures, they often respond with the same adaptive solutions (Conte et al. 2012). For example, parallel changes in gene expression underlying convergent adaptive traits are a well-documented evolutionary phenomenon, with examples from experimental evolution studies imposing uniform selection pressures on replicate populations (Cooper et al. 2003; Riehle et al. 2003), studies in natural systems between closely related taxa (Reid et al. 2016; Derome and Bernatchez 2006; Chan et al. 2010; Nagai et al. 2011; Reed et al. 2011; Manousaki et al. 2013; Zhao et al. 2015), and distantly related taxa (Shapiro et al. 2006; Miller et al. 2007a; Shen et al. 2012). This work has shown that parallelism at the level of gene expression is common in many cases of phenotypic convergence, particularly when divergence time between species is short (Losos 2011; Conte et al. 2012).

However, few studies have investigated the extent of parallel changes in gene expression contributing to species divergence, largely because most expression studies focus on only two species and are either concerned with divergent expression giving rise to divergent phenotypes (Poelstra et al. 2014; Uebbing et al. 2016; Davidson and Balakrishnan 2016) or parallel expression of specific loci (Shapiro et al. 2006; Miller et al. 2007a; Quin et al. 2010) (but see Enard et al. 2002; Ahi et al. 2014). Furthermore, while many genetic and demographic factors are thought to influence the probability of parallel evolution (Rosenblum et al. 2014.; Conte et al. 2012), there are no theoretical expectations for the amount of parallel genetic variation contributing to parallel changes in gene expression during ecological speciation (Schluter et al. 2004; Pavey et al. 2010).

Here we ask whether both parallel and divergent changes in expression underlie novel phenotypes by measuring transcriptomic and genomic divergence between three sympatric species of *Cyprinodon* pupfishes endemic to hypersaline lakes on San Salvador Island, Bahamas. This recent radiation consists of a dietary generalist species (*C. variegatus*) and two novel specialists: a ‘snail-eater’ (*C. brontotheroides*) and a ‘scale-eater’ (*C. desquamator*). Scale-eaters have large jaws and elongated bodies, whereas snail-eaters have short, thick jaws and a protruding nasal region that may function in crushing hard-shelled mollusks. These specialists are more morphologically diverged from one another than either is from their sympatric generalist sister species, and occupy higher trophic levels than the generalist species (Martin and Wainwright 2013a; Martin 2016a, Martin et al. 2017, Hernandez et al. 2017). Scale-eating and snail-eating rapidly evolved within saline lakes on San Salvador Island, Bahamas. These lakes filled within the past 10,000 years after the last glacial maximum (Mylroie and Hagey 1995; Turner et al. 2008), suggesting that speciation occurred rapidly. Scale-eaters and snail-eaters have only been found on San Salvador, and likely diverged from a generalist common ancestor based on phylogenetic analyses of outgroup species (Martin and Wainwright 2011; Martin and Wainwright 2016b). Pupfish populations on many neighboring Bahamian islands and throughout the Caribbean are dietary generalists (Martin and Wainwright 2011; Martin 2016a) and these specialist niches appear unique within atherinomorph and cyprinodontiform fishes (Martin and Wainwright 2011). Indeed, the scale-eating pupfish is separated by 168 million years from other scale-eating fishes (Martin and Wainwright 2013b).

We performed total mRNA sequencing to examine gene expression in lab-reared individuals of all three San Salvador pupfish species from different lake populations at two developmental stages. We also searched 42 whole genomes for SNPs unique to each specialist and determined whether fixed variants near differentially expressed genes showed signs of hard selective sweeps (Pavlidis et al. 2013). We found significant parallelism at the level of gene expression in specialists, but did not find evidence for shared fixed variants underlying parallel changes in expression. We tested whether this counterintuitive result of parallel changes in expression between divergent trophic specialists may be due to 1) decreased pleiotropic constraint for genes showing parallelism or that 2) specialists experience parallel selective environments and adapted to higher trophic levels using similar genetic pathways. Finally, we identified genes differentially expressed between generalists and scale-eaters that contain fixed genetic variants within regions that were previously associated with jaw size variation and showed signs of experiencing a recent hard selective sweep (McGirr and Martin 2017). These regions with fixed variants represent promising *cis*-regulatory elements underlying divergent jaw size – the most rapidly diversifying trait in the San Salvador pupfish radiation (Martin and Wainwright 2013b).

## Methods

### Study system and sample collection

Individuals were caught from hypersaline lakes on San Salvador Island, Bahamas using a hand net or seine net in 2011, 2013, and 2015. Whole genome resequencing was performed for wild-caught individuals from a total of nine isolated lakes on San Salvador (Great Lake, Stout’s Lake, Oyster Lake, Little Lake, Crescent Pond, Moon Rock, Mermaid’s Pond, Osprey Lake, and Pigeon Creek). 14 scale-eaters were sampled from six populations; 11 snail-eaters were sampled from four populations; and 13 generalists were sampled from eight populations on San Salvador. Outgroup samples included one *C. laciniatus* from Lake Cunningham, New Providence Island, Bahamas, one *C. bondi* from Etang Saumautre lake in the Dominican Republic, one *C. diabolis* from Devil’s Hole in California, and captive-bred individuals of *C. simus* and *C. maya* from Laguna Chicancanab, Quintana Roo, Mexico. Sampling is further described in (McGirr and Martin 2017; Richards and Martin 2017). Fish were euthanized in an overdose of buffered MS-222 (Finquel, Inc.) following approved protocols from the University of California, Davis Institutional Animal Care and Use Committee (#17455) and University of California, Berkeley Animal Care and Use Committee (AUP-2015-01-7053) and stored in 95-100% ethanol.

### RNA sequencing and alignment

Juvenile pupfish were derived from either F_0_ wild caught or F_1_ lab raised individuals that were held in a common laboratory environment and fed identical diets (Table S1; 25-27° C, 10-15 ppt salinity, pH 8.3). We collected larvae at two developmental stages: 8-10 and 17-20 days post-fertilization (dpf). The variation in sampling time is due to uncertainty in precise spawning times since eggs were fertilized naturally within breeding tanks and collected on the same day or subsequent day following egg laying. However, we sampled hatched larvae in a haphazard manner over multiple spawning intervals and it is unlikely that sampling time varied consistently by species. Larvae were euthanized in an overdose of buffered MS-222, and stored in RNA later (Ambion, Inc.) at 4° C for one day, followed by long-term storage at −20° C for up to one year. We extracted whole-larvae RNA using RNeasy kits (Qiagen) from 15 larvae (8-10 dpf) (Three F_2_ generalists and F_2_ snail-eaters from Crescent Pond, three F_1_ generalists and F_2_ snail-eaters from Little Lake, and three F_1_ scale-eaters from Little Lake; Table S1). We also dissected 14 larvae (17-20 dpf) to isolate tissues from the anterior craniofacial region containing the dentary, angular articular, maxilla, premaxilla, palatine, and associated craniofacial connective tissues using fine-tipped tweezers washed with RNase AWAY (Three F_2_ generalists and F_2_ snail-eaters from Crescent Pond, three F_1_ generalists and F_2_ snail-eaters from Little Lake, and two F_1_ scale-eaters from Little Lake; Table S1).

Libraries were prepared using the KAPA stranded mRNA-seq kit (KAPA Biosystems 2016) at the High Throughput Genomic Sequencing Facility at UNC Chapel Hill. Stranded sequencing on one lane of Illumina 150PE Hiseq4000 resulted in 677 million raw reads. We filtered raw reads using Trim Galore (v. 4.4, Babraham Bioinformatics) to remove Illumina adaptors and low-quality reads (mean Phred score < 20). We mapped these reads to the *Cyprinodon* reference genome using the RNA-seq aligner STAR (v. 2.5 (Dobin et al. 2013)). We used the featureCounts function of the Rsubread package (Liao et al. 2014) requiring paired-end and reverse stranded options to generate read counts across previously annotated features. We assessed mapping and count quality using MultiQC (Ewels et al. 2016).

### Differential expression analyses

We quantified differences in gene expression between all three species at two developmental stages. Our raw counts determined by featureCounts were normalized with DESeq2 (v. 3.5 (Love et al. 2014)) which uses counts to calculate a geometric mean for each gene across samples, divides individual gene counts by this mean, and uses the median of these ratios as a size factor for each sample. Next, we used DESeq2 to perform pairwise tests pooling species across lakes to identify differentially expressed genes between generalists *vs*. snail-eaters and generalists *vs*. scale-eaters at 8-10 dpf and 17-20 dpf (Table. S1). Genes with fewer than two read counts were discarded from all analyses (n = 1,570), along with genes showing low normalized counts at a threshold determined by DESeq2 (Love et al. 2014). Wald tests determined significant differences in expression between species by comparing normalized posterior log fold change estimates and correcting for multiple testing using the Benjamini–Hochberg procedure with a false discovery rate of 0.05 (Benjamini and Hochberg 1995).

We performed two analyses to test whether specialist species exhibited nonrandom patterns of parallel changes in expression relative to their generalist sister species. We used a Fisher’s exact test to determine whether there was a significant overlap between genes that showed differential expression in both comparisons (i.e. genes that were differentially expressed between generalists *vs*. snail-eaters and generalists *vs*. scale-eaters). A gene that was differentially expressed in both comparisons could either show the same direction of expression in specialists or opposite directions of expression. We performed 10,000 permutations sampling from a binomial distribution to estimate the expected number of genes showing shared and opposite directions of expression. Under this null model of gene expression evolution, a strong positive deviation from 50% of genes showing a shared direction of expression in specialists would indicate significant parallel changes in expression.

Our scale-eater sample sizes were lower than generalist and snail-eater samples for each pairwise comparison (see above). We used a down sampling procedure to test whether sample size affected patterns of parallel changes in expression. We analyzed differential expression for generalists *vs*. snail eaters and generalists *vs*. scale-eaters in 1,000 permutations where generalists and snail-eaters were randomly sampled from our full dataset to match scale-eater sample sizes (n = 3 for 8-10 dpf comparisons; n = 2 for 17-20 dpf). Next, we identified the number of genes differentially expressed between generalists *vs*. snail eaters and generalists *vs*. scale-eaters in each permutation and calculated the proportion of those genes that showed the same direction of expression in specialists relative to generalists. A strong positive deviation from 50% of genes showing a shared direction of expression across permutations would indicate that parallel evolution of expression in specialists is robust to variation in sample size.

### Gene ontology enrichment analyses

We performed gene ontology (GO) enrichment analyses for differentially expressed genes using GO Consortium resources available at geneontology.org (Ashburner et al. 2000; GO Consortium 2017). We used BlastP (v. 2.6 (Camacho et al. 2009)) to identify zebrafish protein orthologs with high similarity (*E*-value < 1) to NCBI protein accessions for genes that we identified as differentially expressed between *Cyprinodon* species. Orthology was established using one-way best hits, where a protein sequence in *Cyprinodon* was the best match to a sequence in zebrafish, and reciprocal best blast hits, where a sequence in *Cyprinodon* was the best match to a sequence in zebrafish and vice versa. While reciprocal best hits robustly predict orthology with high precision, it is highly conservative and fails to detect many true orthologs in duplication rich clades such as teleosts (Altenhoff and Dessimoz 2009; Salichos and Rokas 2011; Dalquen and Dessimoz 2013). Thus, we performed GO enrichment analyses using orthologs defined as one-way best hits, and compare these results to enrichment analyses using more conservative orthologs defined as reciprocal best hits.

Genes were either differentially expressed between generalists and snail-eaters, generalists and scale-eaters, or in both comparisons. Thus, we performed two GO enrichment analyses for: 1) genes that were differentially expressed in both companions, and 2) genes differentially expressed in one comparison. We grouped enriched GO categories into similar representative terms using the REVIGO clustering algorithm (Tomislav 2011). REVIGO groups semantically similar terms to reduce the size and redundancy of lists from GO enrichment analyses, where grouping is guided by *P*-values corrected for multiple comparisons (Tomislav 2011). When similar terms show similar enrichment, they are assigned to a single representative term. We measured differences in the proportion of representative terms describing metabolic and developmental processes between genes showing parallel and divergent changes in expression between specialists.

### Measuring pleiotropy for differentially expressed genes

The probability of parallel evolution of gene expression may be higher for genes that are less constrained by pleiotropy (Cooper and Lenski 2000; Manceau et al. 2010; Rosenblum et al. 2014). High gene pleiotropy is correlated with participation in more protein-protein interactions (PPIs), which in turn effects multiple biological processes (He and Zhang 2006; Safari-alighiarloo et al. 2014). Genes that act across multiple developmental stages are also more pleiotropic (Stern and Orgogozo 2008). We used one-way best hits zebrafish orthologs to estimate pleiotropy for differentially expressed genes based on their number of associated GO biological processes, PPIs, and developmental stages when they are known to be expressed (Papakostas et al. 2014). We again used GO Consortium resources (Ashburner et al. 2000; GO Consortium 2017) to determine the number of biological processes associated with each gene. We examined biological process annotations only for genes from ZFIN (zfin.org) with experimental evidence (GO evidence code EXP). The String protein database (v. 10; (Szklarczyk et al. 2015)) calculates a combined score measuring confidence in protein interactions by considering known interactions (experimentally determined and from manually curated databases) and predicted interactions. We used the String database to quantify PPIs for protein products of differentially expressed genes, focusing only on interactions with experimental evidence (i.e. non-zero experimental evidence scores). Next, we determined the number of developmental stages where a gene is known to be expressed using the Bgee expression call database for zebrafish (v. 14.0 (Bastian et al. 2008)). We considered eight developmental stages from larval day five to juvenile day 89 from the Zebrafish Stage Ontology (ZFS) that were deemed ‘gold quality,’ meaning there was no contradicting call of absence of expression for the same gene, in the same developmental stage (Bastian et al. 2008).

We tested whether genes showing parallel changes in expression between specialists showed lower levels of pleiotropy than genes showing divergent changes in expression by fitting a generalized linear model on count data for pleiotropy estimates (negative binomial family; *glm.nb* function in the R library “MASS”). We did not measure pleiotropy for genes expressed at 17-20 dpf due to the low number of zebrafish orthologs matched for genes with parallel expression in craniofacial tissues (11 out of 23).

### Genomic variant discovery and population genetic analyses

SNP variants were called using previously outlined methods (McGirr and Martin 2017; Richards and Martin 2017). Briefly, 42 individual DNA samples extracted from muscle tissue were fragmented, barcoded with Illumina indices, and quality checked using a Fragment Analyzer (Advanced Analytical Technologies, Inc.). Sequencing on four lanes of Illumina 150PE Hiseq4000 resulted in 2.8 billion raw reads that were mapped from 42 individuals to the *Cyprinodon* reference genome (NCBI, *C. variegatus* Annotation Release 100, total sequence length = 1,035,184,475; number of scaffold = 9,259, scaffold N50, = 835,301; contig N50 = 20,803; (Lencer et al. 2017)). We followed Genome Analysis Toolkit (v 3.5) best practices and hard filter criteria to call and refine our SNP variant dataset (QD < 2.0; FS < 60; MQRankSum < −12.5; ReadPosRankSum < −8 (DePristo et al. 2011)). We filtered our final SNP dataset to include individuals with a genotyping rate above 90% (no individuals were excluded by this filter) and SNPs with minor allele frequencies higher than 5%, resulting in 16 million variants with a mean sequencing coverage of 7× per individual (range: 5.2–9.3×).

We identified SNPs that were fixed in each specialist species. We calculated genome wide F_st_ using VCFtools’ ‘weir-fst-pop’ function for two different population comparisons involving samples collected from San Salvador: generalists (n = 13) *vs*. snail-eaters (n = 11) and generalists (n = 13) *vs*. scale-eaters (n = 9). Differences in sample sizes made our analyses biased to detect more fixed variation between generalists vs. scale-eaters (n = 13 vs. 9) than between generalists vs. snail-eaters (n = 13 vs. 11). We also performed 1,000 permutations calculating genome wide F_st_ between randomly subsampled groups in order to identify non-randomly differentiated genomic regions between species. We calculated the 99th percentile estimates of F_st_ across all SNPs between randomly sampled generalists and snail-eaters (n = 13 vs. n = 11) and between randomly sampled generalists and scale-eaters (n = 13 vs. n = 9). We took the 99th percentile of these distributions to set a threshold defining significantly divergent outliers (Fig. S7).

Our SNP dataset included 14 scale-eaters, however, we split our scale-eater population into two groups (large-jawed scale-eaters, n = 9 and small-jawed scale-eaters, n = 5) based on previous evidence that these two populations are genetically distinct (McGirr and Martin 2017; Richards and Martin 2017). This allowed us to identify SNPs unique to large-jawed scale-eaters (i.e. *C. desquamator* (Martin and Wainwright 2013a)), which were the only type of scale-eater we sampled for RNA-seq. We identified which of these SNPs resided in gene regions (either exonic, intronic, or within 10kb of the first or last exon) for genes showing differential expression. We determined whether these regions showed signatures of hard selective sweeps using SweeD ((Pavlidis et al. 2013); methods previously described in (McGirr and Martin 2017)). Briefly, SweeD sections scaffolds into 1,000 windows of equal size and calculates a composite likelihood ratio (CLR) using a null model where the site frequency spectrum of each window does not differ from that of the entire scaffold. We previously estimated ancestral effective population sizes of San Salvador pupfishes using MSMC (Schiffels and Durbin 2014; McGirr and Martin 2017) and used these estimates to correct the expected neutral site frequency spectrum for the inferred recent population bottleneck in Caribbean pupfishes using SweeD. Windows with fixed SNPs that showed CLRs above the 95^th^ percentile across their respective scaffolds (>10,000bp) under the assumptions of a recent population bottleneck were interpreted as regions that recently experienced a hard sweep.

## Results

### Differential expression between generalists and each specialist

Total mRNA sequencing across all 29 samples resulted in 677 million raw reads, which was reduced to 674 million reads after quality control and filtering. 81.2% of these reads successfully aligned to the reference genome and 75.5% of aligned reads mapped to annotated features with an average read depth of 309× per sample. The number of reads mapping to annotated features was comparable across generalists, snail-eaters, and scale-eaters (ANOVA; 8-10 dpf *P* = 0.47; 17-20 dpf *P* = 0.33; Fig. S1).

Snail-eaters and scale-eaters occupy novel niches among over 2,000 species of atherinomorph fishes (Martin and Wainwright 2011), and these trophic specialist species likely evolved from a generalist common ancestor within the past 10,000 years (Mylroie, J.E, Hagey 1995; Turner et al. 2008). We analyzed transcriptomic changes underlying rapid trophic divergence by comparing specialist species gene expression against their sympatric generalist sister species. We used DESeq2 to identify genes that were differentially expressed between generalists *vs*. snail-eaters and generalists *vs*. scale-eaters at 8-10 days post-fertilization (whole body tissue) and 17-20 dpf (craniofacial tissue).We measured expression across 22,183 genes with greater than two read counts out of 24,383 total genes annotated for the *Cyprinodon variegatus* assembly (NCBI, *C. variegatus* Annotation Release 100, (Lencer et al. 2017)).

At 8-10 dpf, we found 1,014 genes differentially expressed between generalists *vs*. snail-eaters and 5,982 genes differentially expressed between generalists *vs*. scale-eaters (Fig. 1A and C; Fig. 2A) (Benjamini and Hochberg adjusted *P* ≤ 0.05). 818 genes were differentially expressed in both comparisons, which is a significantly larger amount of overlap than expected by chance (Fisher’s exact test, *P* < 1.0 × 10^−16^). Remarkably, 815 of these 818 genes showed the same direction of expression in specialists relative to generalists (Fig. 2B). Specifically, 441 differentially expressed genes showed lower expression in both specialist species compared to generalists, while 374 showed higher expression in specialists. Only three genes showed opposite directions of expression (Fig. 2B). Two genes showed higher expression in snail-eaters and lower expression in scale-eaters while one gene showed higher expression in scale-eaters (Table S2). This is significantly more parallel change in expression between specialists than would be expected under a null model of gene expression evolution, where a gene has an equal chance of showing a shared or opposite direction of expression in specialists relative to generalists (10,000 permutations, *P* < 1.0 × 10^−4^; Fig. S2). Parallel evolution of expression in specialists was consistent at both the gene and isoform level (Fig. S2, S3).

**Fig. 1.**
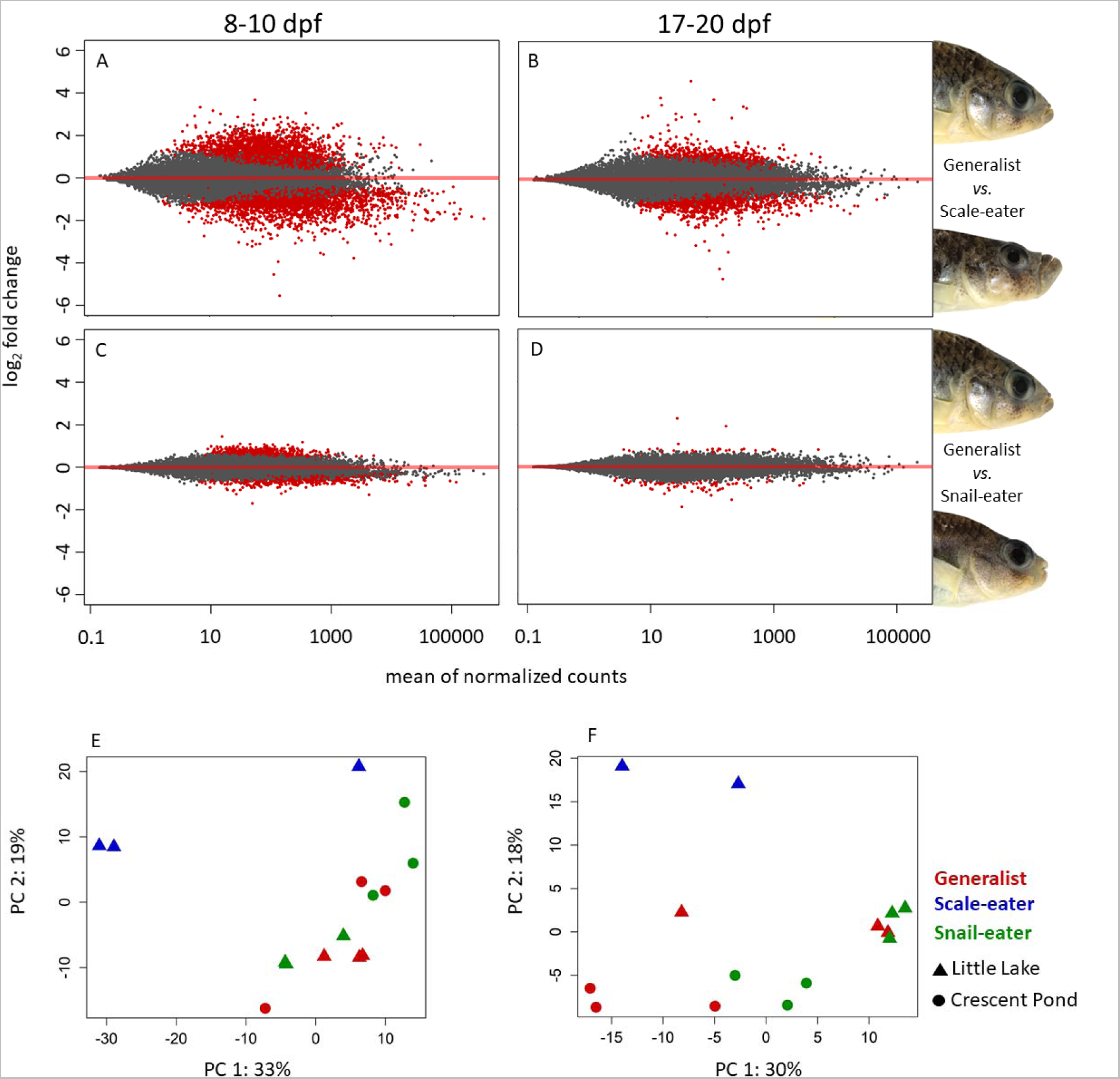
Differential gene expression between generalists and trophic specialists. Red points represent genes that are differentially expressed in 8-10 dpf whole-larvae tissue (A, C) and 17-20 dpf craniofacial tissue (B, C) between generalists *vs*. scale-eaters (A, B) and generalist *vs*. snail-eaters (C, D). Bottom panels show the top two principal components accounting for a combined 52% (8-10 dpf; E) and 48% (17-20 dpf; F) of the total variation between samples across 413 million reads mapped to annotated features. Triangles represent samples from Little Lake and circles represent samples from Crescent Pond on San Salvador Island.

**Fig. 2.**
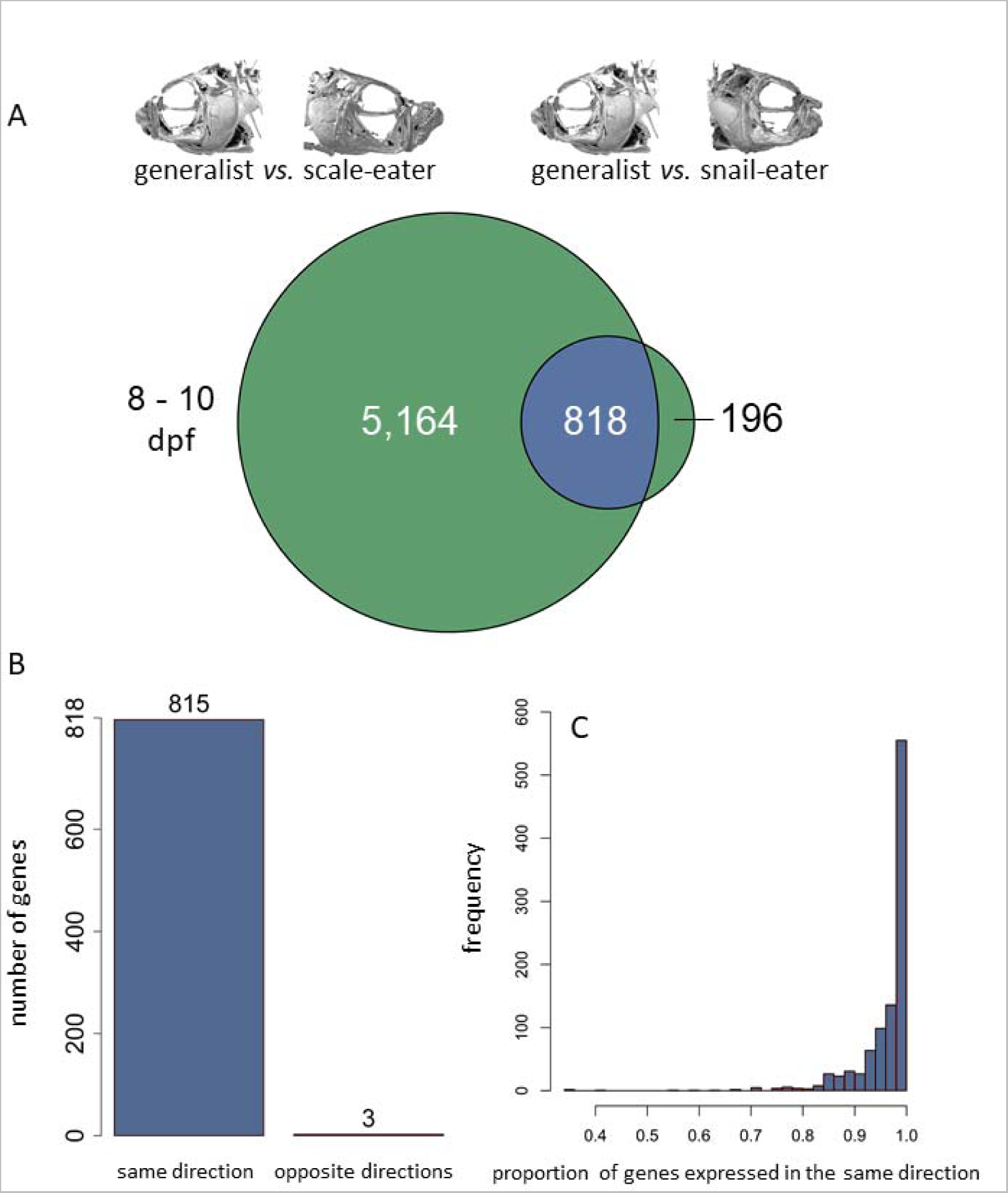
Parallel evolution of gene expression between specialists despite divergent trophic adaptation. A) Circles illustrate genes differentially expressed in 8-10 dpf whole-larvae tissue for generalists *vs*. scale-eaters (left) and generalists *vs*. snail-eaters (right). Genes showing differential expression in both comparisons are shown in blue, and those showing divergent expression patterns unique to each specialist are green. Significantly more genes show differential expression in both comparisons than expected by chance (Fisher’s exact test, *P* < 1.0 × 10^−16^). B) Significantly more genes show the same direction of expression in specialists relative to generalists than expected by chance (10,000 permutations; *P* < 1.0 × 10^−4^; Fig. S2). C) Distribution of the proportion of genes differentially expressed in the same direction between specialists relative to after 1,000 down sampling permutations show that parallel expression is robust to variation in sample size (median number of genes common to both comparisons = 61).

Craniofacial morphology is the most rapidly diversifying phenotypic axis measured so far within the San Salvador radiation (Martin and Wainwright 2013b; Martin 2016a). In order to detect genes expressed during jaw development, we compared expression within craniofacial tissue at the 17-20 dpf stage. We found a similar pattern of parallel changes in gene expression at this developmental stage (Fig. S4). 120 genes were differentially expressed between generalists *vs*. snail-eaters and 1,903 genes differentially expressed between generalists *vs*. scale-eaters (Fig. 1B and D). Again, we saw a significant amount of overlap between comparisons with 23 genes differentially expressed in both comparisons (Fisher’s exact test, *P* < 1.0 × 10^−5^). 22 of these 23 genes showed the same direction of expression in specialists relative to generalists (Fig. S4). Specifically, 10 genes showed lower expression in both specialist species compared to generalists, while 12 showed higher expression in specialists (Fig. S4). Only one gene (*mybpc2*) showed opposite directions of expression, with higher expression in snail-eaters and lower expression in scale-eaters (Table S2).

Our sample sizes for scale-eater species were lower for comparisons at 8-10 dpf (n = 3) and 17-20 dpf (n = 2) relative to snail-eaters and generalists (n = 6 for both stages). We measured differential expression for generalists *vs*. snail eaters and generalists *vs*. scale-eaters in 1,000 permutations where we randomly down-sampled generalists and snail-eaters from our full dataset to match scale-eater sample sizes. Figure 2C shows the proportion of genes differentially expressed at 8-10 dpf in both comparisons that showed the same direction of expression in specialists relative to generalists across 1,000 permutations. The total number of differentially expressed genes in each permutation was variable (Fig. S5A and C, median number of genes common to both comparisons = 61). Despite this variability, we found that the parallel evolution of expression in specialists was robust to smaller sample size, with greater than 90% of genes showing parallel evolution of expression in 90% of permutations (Fig. 2C). However, at 17-20 dpf parallel changes in expression were not as consistent across permutations (Fig. S4C and S5F).

### Genes showing parallel changes in expression are enriched for metabolic processes

We performed GO enrichment analyses with one-way blast hit zebrafish orthologs for genes showing parallel changes in expression between specialists (n = 620) and genes showing divergent expression patterns in snail-eaters (n = 102) and scale-eaters (n = 3,349). We restricted these analyses to genes expressed at 8-10 dpf because the number of genes showing parallel expression in specialists at 17-20 dpf (n = 23) was low and did not show enrichment for any biological process.

We grouped enriched GO categories into similar representative terms using the REVIGO clustering algorithm (Tomislav 2011). Genes showing parallel changes in expression between specialists were enriched for metabolic processes (20% of representative terms; Fig. 3A; Table S3). In contrast, genes with divergent expression patterns in specialists were enriched for cranial skeletal development and pigment biosynthesis (7% and 3% of terms, respectively) while only 11% of enriched categories described metabolic processes (Table S4).

**Fig. 3.**
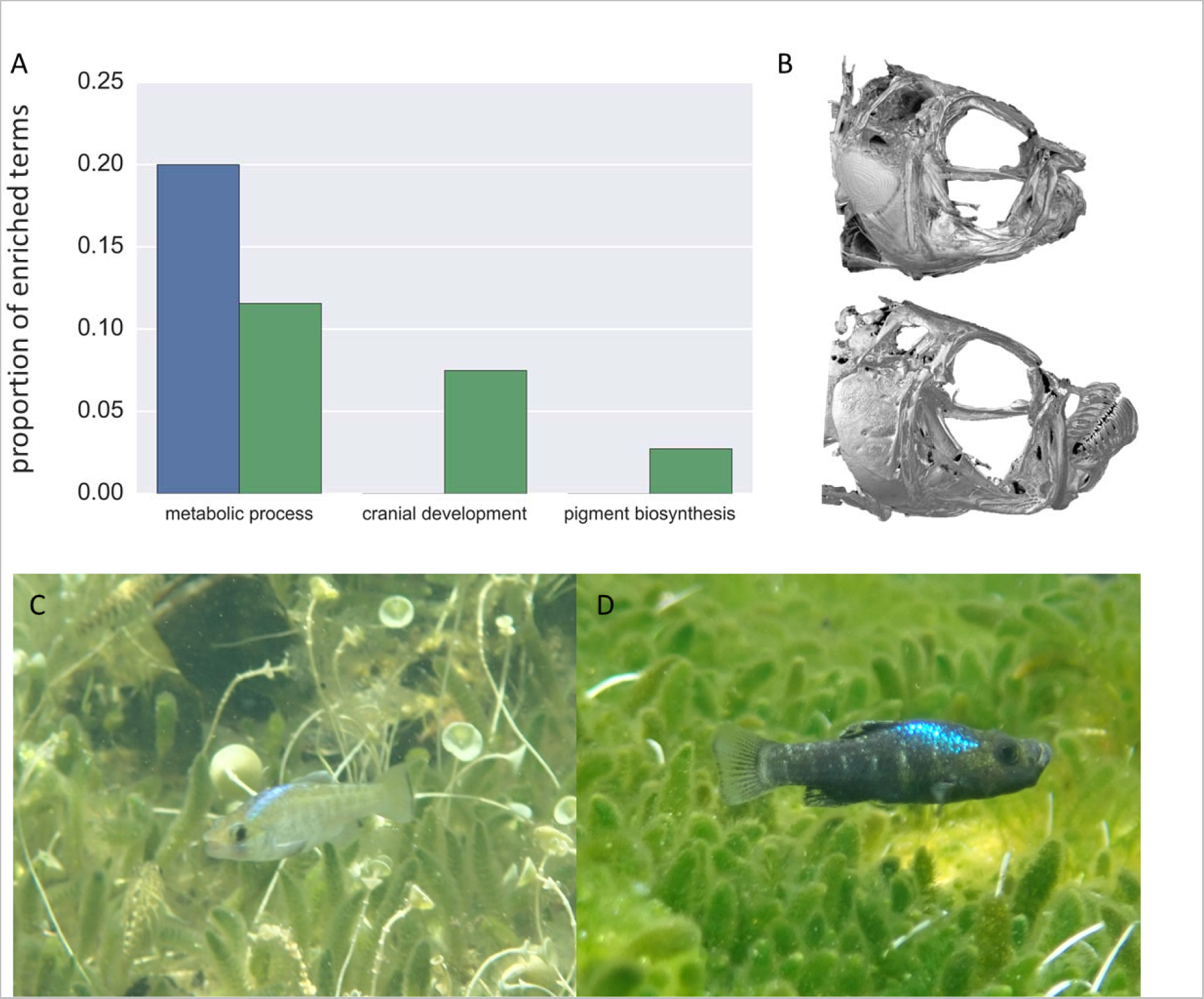
Parallel gene expression underlies metabolic adaptations while divergent expression underlies trophic morphology. A) Genes showing parallel changes in expression between specialists (blue) and genes showing divergent expression (green) are contrastingly enriched for terms describing metabolic processes (parallel: 20% of enriched terms; divergent: 11% of terms). Genes showing divergent expression are enriched for cranial skeleton development (7% of terms) and pigment biosynthesis (3% of terms). B) μCT scans show drastic craniofacial divergence between snail-eaters (top) and scale-eaters (bottom) (modified from Hernandez et al. 2017). Bottom panels show male breeding coloration characteristic of light snail-eaters (C) and dark scale-eaters (D).

We also performed GO enrichment analyses using orthologs that were established using a more conservative reciprocal best hit approach, where a sequence in *Cyprinodon* was the best match to a sequence in zebrafish and vice versa. As expected, we identified fewer reciprocal best hits than one-way hits (615 genes showing parallel changes in expression between specialists, 95 genes showing divergent expression unique to snail-eaters, and 2,150 genes showing divergent expression unique to scale-eaters). Encouragingly, we still found that genes showing parallel changes in expression were enriched for metabolic processes (26% of representative terms), whereas genes showing divergent expression showed less enrichment for metabolic processes (20% of representative terms). However, we did not see any enrichment for cranial development or pigment biosynthesis for genes showing divergent expression using reciprocal best hit orthologs.

We tested whether genes showing parallel changes in expression were less constrained by pleiotropy than genes showing divergent expression between specialists. We estimated pleiotropy for orthologs of differentially expressed genes based on their number of protein-protein interactions (PPIs), associated GO biological processes, and developmental stages when they are known to be expressed. However, we did not find any difference in pleiotropy for genes showing parallel changes in expression compared to genes showing divergent expression using any of these three metrics (GLM; biological processes: *P* = 0.67; PPIs: *P* = 0.09; developmental stages: *P* = 0.89) (Fig. S6).

### Genetic variation underlying parallel changes in expression

We identified 79 SNPs fixed between generalists *vs*. snail-eaters and 1,543 SNPs fixed between generalists *vs*. scale-eaters (also see our previous study on genome-wide association mapping jaw length in these species). None of these fixed variants were shared between specialists. Next, we determined which of these fixed SNPs fell within gene regions (either exonic, intronic, or within 10kb of the first or last exon; Table 1). 26 SNPs fixed in snail-eaters overlapped with 17 gene regions, whereas 1,276 SNPs fixed in scale-eaters overlapped with 245 gene regions.

**Table 1.** Genomic distribution of fixed variants. The first five columns show the total number of fixed SNPs in each species comparison and how many fall within exons, introns, 10kb of the first or last exon of a gene, and outside of 10kb from the first or last exon of a gene. Final two columns show the number of genes with fixed SNPs within the gene and/or within 10kb of the first or last exon. The last column shows the number of differentially expressed (DE) genes near fixed SNPs that includes DE genes from 8-10 dpf and 17-20 dpf comparisons.

| comparison | fixed SNPs | exonic | intronic | within 10kb<br>of coding<br>region | >10kb away<br>from coding<br>region | number of<br>genes near<br>fixed SNPs | number of<br>DE genes near<br>fixed SNPs |
| --- | --- | --- | --- | --- | --- | --- | --- |
| generalist<br>vs.<br>snail-eater | 79 | 0 | 16 | 10 | 53 | 17 | 1 |
| generalist<br>vs.<br>scale-eater | 1,543 | 140 | 512 | 624 | 267 | 245 | 83 |

Next, we identified fixed variants near genes that showed differential expression. We found 319 SNPs fixed in scale-eaters within 71 gene regions that showed differential expression between generalists and scale-eaters at 8-10 dpf and 118 SNPs within 26 gene regions differentially expressed between generalists and scale-eaters at 17-20 dpf. We suspect that some of these fixed variants are within *cis*-regulatory elements responsible for species-specific expression patterns that ultimately give rise to phenotypic differences in scale-eaters. Conversely, we only identified a single SNP fixed in snail-eaters within a gene (*tmprss2*) that was differentially expressed between generalists and snail-eaters at 8-10 dpf. We did not find any fixed variants near genes differentially expressed between generalists and snail-eaters at 17-20 dpf, possibly suggesting that fixed variants regulate expression divergence at an earlier developmental stage.

Since we did not find any variants that were fixed between snail-eaters and generalists that were also fixed between scale-eaters and generalists, we searched for shared variation at a lower threshold of genetic divergence. We calculated the 99^th^ percentile outlier F_st_ estimates between randomly subsampled groups of each species across 1,000 permutations to create two null distributions of genome-wide divergence. We took the 99^th^ percentile of these distributions as an estimate of significantly high divergence (Fst > 0.36 for generalists *vs*. snail-eaters; Fst > 0.42 for generalists *vs*. scale-eaters; Fig. S7). We found 4,410 SNPs above this lower threshold of divergence near 134 genes showing parallel changes in expression between specialists at 8-10 dpf. The most differentiated SNPs near genes showing parallel changes in expression show F_st_ < 0.8 between generalists *vs*. snail-eaters and generalists *vs*. scale-eaters. Overall, these results suggest it is unlikely that the parallel evolution of gene expression in specialists is controlled by shared variation that is fixed or nearly fixed in specialist populations.

### The genetic basis of extreme craniofacial divergence

We previously described 30 candidate gene regions containing variants fixed between trophic specialist species associated with variation in jaw length. These candidates also showed signatures of a recent hard selective sweep (McGirr and Martin 2017). Encouragingly, we found ten of these genes differentially expressed between generalists and scale-eaters (eight at 8-10 dpf and two at 17-20 dpf) and one between generalists and snail-eaters (8-10 dpf; Table S5).

We searched for signatures of hard selective sweeps across the 84 gene regions containing fixed variation in specialists (Table 1). Interestingly, 80% of these gene regions showed signs of a hard sweep (estimated by SweeD; CLR > 95^th^ percentile across their respective scaffolds; Table S6). All of these gene regions contained SNPs that were either fixed between generalists *vs*. snail-eaters or generalists *vs*. scale-eaters and showed differential expression at 8-10 dpf, 17-20 dpf, or both. Finally, we compared this list of genes experiencing selection to those annotated for cranial skeletal system development (GO:1904888) and muscle organ development (GO:0007517). While this search was limited to zebrafish orthologs identified as one-way best hits, we were able to identify three genes containing fixed variation in scale-eaters that likely influence craniofacial divergence through cis-acting regulatory mechanisms (*loxl3b* (annotated for cranial effects); *fbxo32* and *klhl40a* (annotated for muscle effects)).

## Discussion

We combined RNA sequencing with genome-wide divergence scans to study the molecular evolution of two trophic specialist species that rapidly diverged from a generalist common ancestor within the last 10,000 years. We examined how gene expression and SNP variation influence snail-eater and scale-eater niche adaptations using comparisons between each specialist and their generalist sister species. We found a significant amount of parallelism at the level of gene expression yet no parallelism at the level of fixed genetic variation within specialists. Specifically, 80% of genes that were differentially expressed between snail-eaters and generalists were up or downregulated in the same direction when comparing expression between scale-eaters and generalists (Fig. 2A). We explored two possible explanations for this pattern: 1) reduced pleiotropic constraints made these genes likely targets for parallelism or 2) convergent processes drove parallel gene expression evolution in this highly divergent pair of specialist species due to shared adaptations to a higher trophic level.

### 1. Pleiotropic constraints do not explain parallel changes in gene expression

Genes that effect one or a few traits are less constrained than genes with many phenotypic effects, perhaps making them simpler shared targets for expression divergence during adaptive evolution between independently evolving lineages. Indeed, theory predicts that the probability of parallel evolution of gene expression should be higher for genes with minimal pleiotropic effects (Manceau et al. 2010, Rosenblum et al. 2014). We predicted that genes showing parallel changes in expression between specialists would show lower degrees of pleiotropy than divergently expressed genes. We estimated three measures of gene pleiotropy (number of associated GO biological processes, protein-protein interactions (PPIs), and developmental stages when they are known to be expressed) and found no significant difference in any measure for genes showing parallel versus divergent changes in expression patterns (Fig. S6). This finding is consistent with some empirical evidence and theoretical models of gene expression evolution that found pleiotropy constrains the variability of gene expression within species, but does not hinder divergence between species (Tulchinsky et al. 2014; Uebbing et al. 2016).

### 2. Parallel changes in gene expression underlie convergent metabolic adaptations to a higher trophic level in each specialist

While the specialists are more morphologically diverged from one another than either is from the generalist species, particularly in their craniofacial phenotype and male reproductive coloration (Martin and Wainwright 2013a; Martin et al. 2017) (Fig. 3B and C), dietary isotope analyses show that they both occupy a higher trophic level than generalists (Martin 2016b). Fish scales and mollusks contribute to more nitrogen-rich diets in specialists compared to generalist species that primarily consume algae and detritus (Martin 2016b). Perhaps the same metabolic processes required for this type of diet are adaptive at higher trophic levels for both scale-eaters and snail-eaters, which might explain patterns of parallel changes in expression. Thus, we predicted that genes showing parallel changes in expression would affect metabolic processes that may be similar between specialists, whereas genes showing divergent expression between specialists would affect morphological development.

GO enrichment analyses using one-way best-hit zebrafish orthologs support both hypotheses. We found that 20% of GO terms enriched for genes showing parallel changes in expression described metabolic processes, and zero described cranial skeletal development or pigment biosynthesis (Fig. 3A; Table S3). In contrast, 10% of terms showing enrichment in the divergently expressed gene set described developmental processes (cranial skeletal development and pigment biosynthesis) and only 11% described metabolic processes (Fig 3A, Table S4). GO enrichment analyses using more conservatively defined reciprocal best hit orthologs confirmed that genes showing parallel changes in expression were highly enriched for metabolic processes (26% of representative terms). These results suggest that the parallel evolution of expression in specialists confers adaptation to a higher trophic level. Snail-eating and scale-eating may present similar metabolic requirements relative to the lower trophic level of algivorous generalists. This is consistent with the high macroalgae content of generalist diets relative to both specialist species (Martin and Wainwright 2013b) and the shorter intestinal lengths observed in both specialists relative to the generalist (CHM and JAM personal observation).

Enrichment analyses using one-way best hit orthologs indicate that genes showing divergent expression in specialists are responsible for shaping divergent cranial and pigmentation phenotypes between species (Fig. 3), but we did not find enrichment for these processes using reciprocal best hit orthologs. This may be because up to 60% of orthologous relationships are missed by the reciprocal best-hit criterion in lineages with genome duplications, including teleosts (Dalquen and Dessimoz 2013). Finally, both approaches we used to establish orthology indicated that genes showing divergent expression in specialists were moderately enriched for metabolic processes (Fig. 3A; Table S4). While parallel changes in expression may broadly influence adaptation to a higher trophic level, these divergently expressed metabolic genes likely play a role in dietary specialization unique to each species.

### Parallel changes in gene expression despite unshared genetic variation

We find significant parallel evolution of gene expression across genes that are annotated for effects on metabolism, yet shared expression patterns do not seem to be driven by the same fixed variants. This is surprising in this young radiation given that the probability of shared genetic variation underlying phenotypic convergence increases with decreasing divergence time (Schluter et al. 2004; Conte et al. 2012; Martin and Orgogozo 2013). Although 80% of differentially expressed gene regions containing fixed SNPs show signs of experiencing a selective sweep, and almost none of these variants were in exons, it is still possible that fixed alleles do not regulate parallel changes in expression for metabolic genes. Indeed, we found 4,410 SNPs that showed significant differentiation between generalists *vs*. snail-eaters and generalists *vs*. scale-eaters near 134 genes showing parallel changes in expression. These shared variants all showed F_st_ < 0.8, suggesting that parallel expression is not controlled by shared variation that is fixed or nearly fixed in specialist populations. However, our results do not rule out a role for fixed variation influencing the parallel evolution of expression through long-range chromosome interactions or during earlier critical developmental stages, such as neural crest cell migration at approximately 48 hpf.

Many studies that show parallel adaptation at the gene level describe convergence within pigmentation and skeletal development pathways (Miller et al. 2007b; Reed et al. 2011; Conte et al. 2012; Kronforst et al. 2012). Perhaps the architecture of metabolic adaptation is more flexible, having more mutational targets or employing more late-acting developmental regulatory networks that are less constrained than early-acting networks (Kalinka et al. 2010; Garfield et al. 2013; Martin and Orgogozo 2013; Reddiex et al. 2013; Ferna et al. 2014; Comeault et al. 2017). Our findings highlight the importance of understanding convergence across different biological levels of organization.

### Candidate genes influencing trophic adaptations

We found many genes affecting metabolism that were differentially expressed in the same direction in specialists relative to their generalist sister species. While the metabolism ontology includes a broad class of proteins with a variety of biological functions, we find many with distinct effects on dietary metabolism. For example, the gene *asl* (argininosuccinate lyase) is important for nitrogen excretion. Variants of *asl* are associated with argininosuccinic aciduria and citrullinemia, conditions involving an accumulation of ammonia in the blood (Saheki et al. 1987; Hu et al. 2015). This gene, along with some of 274 other genes we found annotated for nitrogen metabolism, may show parallel changes in expression between specialists as an adaptation to nitrogen-rich diets (Martin 2016b).

We also identified candidate genes influencing cranial divergence that were differentially expressed between scale-eaters and generalists, contain SNPs fixed in scale-eaters, and showed signs of a hard selective sweep. *loxl3b* is highly expressed in scale-eaters at 8-10 dpf and annotated for cranial effects (Table S6). The protein encoded by this gene (lysyl oxidase 3b) controls the formation of crosslinks in collagens, and is vital to cartilage maturation during zebrafish craniofacial development (Van Boxtel et al. 2011). Mutations in *loxl3b* are associated with Stickler Syndrome, which is characterized by cranial anomalies and cleft palate (Alzahrani et al. 2015). *fbxo32* and *klhl40a* are both expressed at lower levels in scale-eaters at 8-10 dpf relative to generalists and may influence skeletal muscle divergence between species (Table S6). High expression of *fbxo32* is associated with muscle atrophy, while mutations in *klhl40a* cause nemaline myopathy (muscle weakness) (Ravenscroft et al. 2013; Mei et al. 2015). Variants fixed in scale-eaters near these genes, along with fixed variation near differentially expressed genes previously associated with large jaw size (McGirr and Martin 2017; Table S5) represent strong *cis*-acting regulatory candidates potentially influencing scale-eater cranial traits.

### Caveats to gene expression analyses and the robustness of parallel evolution

We compared the transcriptomes of derived trophic specialists to a contemporary generalist sister species to identify gene expression divergence important for the evolution of trophic traits. However, the generalist transcriptome represents an approximation of the putative ancestral state, and has also evolved over the past 10,000 years (Holtmeier 2001; Turner et al. 2008; Martin and Wainwright 2011; Martin 2016a). We chose to sample RNA at 8-10 dpf and 17-20 dpf to identify transcriptional variation that influences larval development, however, some activation of parallel gene networks is likely specified at pre-hatching developmental stages (Garfield et al. 2013; Ferna et al. 2014). It is also possible that we did not have the power to identify subtle differences in expression for genes that showed high divergence between specialists and generalists. Detecting differential expression of transcripts is notoriously difficult when read counts are low and variance within treatment groups is high (Conesa et al. 2016; Lin et al. 2016). We were able to detect differential expression for genes with a mean normalized count as low as 1.6 (median = 150) and log_2_ fold change as low as 0.2 (median = 1.11). Furthermore, our scale-eater sample sizes (8-10 dpf n = 3; 17-20 dpf n = 2) were lower than that of generalists and snail-eaters (n = 6 at both stages; Table S1). Nonetheless, down sampling analyses suggest that patterns of parallel expression are robust to smaller sample sizes for 8-10 dpf tissue (Fig. 2C), but less so for 17-20 dpf tissue (Fig. S4C).

Finally, our novel results are consistent with a recently published independent analysis of gene expression in San Salvador pupfishes that identified many of the same genes we found divergently expressed between specialists (Lencer et al. 2017). We examined this dataset using the same significance thresholds for differentially expressed genes as described in Lencer *et al.* for mRNA extracted from all three species at 8 dpf and 15 dpf (*P* < 0.1 and |Log_2_ fold change| > 0.2). We found that 40% of genes divergently expressed between specialists in this dataset were divergently expressed in our own dataset. Importantly, Lencer *et al.* only examined cranial tissues at both of these developmental stages and they did not choose to examine parallel evolution of expression. We also searched for evidence of parallel change in expression for mRNA extracted from all three species at 8 dpf in the Lencer et al. dataset. 28.8% of genes that were differentially expressed between snail-eaters and generalists were up or downregulated in the same direction between scale-eaters and generalists. This is a lower proportion of parallel change in expression than we identified (Fig. 2), but this is most likely because Lencer *et al.* only sampled RNA from cranial tissues at 8 dpf, unlike our sampling of whole larvae. Thus, the majority of parallel changes in expression between specialists likely occurs in non-cranial tissues, consistent with our shared metabolic hypothesis.

## Conclusion

Here we find significant parallel evolution of gene expression between two highly divergent specialist species relative to their generalist sister species. While there are many cases of parallel changes in expression underlying parallel specialization, to our knowledge, this represents the first case of parallel expression underlying divergent specialization. Numerous studies have shown that shared genetic variation underlying phenotypic convergence is more likely when divergence times between species are short (Schluter et al. 2004; Conte et al. 2012; Martin and Orgogozo 2013). Scale-eating and snail-eating species have evolved rapidly within the last 10,000 years, yet we do not find the same variants fixed in both species underlying parallel changes in expression. We show that parallel evolution of expression likely reveals convergent adaptation to a higher trophic level in each specialist, despite their highly divergent resource use and morphology.

## Acknowledgements

This study was funded by the University of North Carolina at Chapel Hill and the Miller Institute for Basic Research in the Sciences to CHM. We thank Sara Suzuki, Jelmer Poelstra, and Emilie Richards for valuable discussions and computational assistance; the High Throughput Genomic Sequencing Facility at UNC Chapel Hill for performing RNA library prep and Illumina sequencing; the Gerace Research Centre for accommodation; and the Bahamian government BEST Commission for permission to conduct this research. All datasets used for this study will be deposited in the NCBI Short Read Archive associated with BioProject PRJNA391309.

## Author Contributions

JAM wrote the manuscript. JAM extracted the RNA samples, and conducted all bioinformatic and population genetic analyses. Both authors contributed to the conception and development of the ideas and revision of the manuscript.

## Data Accessibility

All genomic and transcriptomic raw sequence reads are available on the NCBI BioProject database. Title: Craniofacial divergence in Caribbean Pupfishes. Accession: PRJNA391309.

## Competing interests

We declare we have no competing interests.

